# Impaired Sensory-motor Reconfiguration and Pupil-linked Arousal in Aging

**DOI:** 10.64898/2026.06.01.729265

**Authors:** Maria J. Ribeiro, Joshua Calder-Travis, Nádia Canário, Miguel Castelo-Branco, Tobias H. Donner

## Abstract

Mounting evidence points to a key role of the brainstem arousal systems in controlling the adaptation of behavior to changing contexts. Both, brainstem arousal systems and cognitive flexibility seem to deteriorate in aging. We hypothesized that these two impairments are related. To test this hypothesis in humans, we compared adaptive behavior and arousal dynamics between older and younger adults (60% women). Arousal dynamics were measured by means of luminance-independent pupil responses. Participants performed a hierarchical decision-making task, which required frequent switches between two stimulus-response mapping rules for reporting a perceptual categorization judgment. These rule switches were hidden, and participants needed to infer them from noisy evidence. Older adults were less accurate than younger adults. This impairment was due to more frequent use of the incorrect rule. Neither the perceptual categorization nor the detection of rule switches differed between age groups. Moments of high probability of rule switches were associated with pupil dilation in both age groups, which was, however, reduced in older people. We propose that impaired cognitive recruitment of pupil-linked arousal in aging might limit the ongoing reconfiguration of the neural pathways that support flexible cognitive behavior.

**Significance Statement:** Aging affects cognitive flexibility, but the physiological basis of these alterations is poorly understood. Theoretical work and mounting empirical evidence indicate that the ascending neuromodulatory systems of the brainstem may play a key role in the dynamic reconfiguration of functional brain networks underlying cognitive flexibility. Here, we show an impaired performance of older compared to younger adults in a decision-making task that requires frequent switching between different stimulus-response associations. Critically, older adults also exhibited an abnormal recruitment of their brainstem arousal systems triggered by the switches in the task. We propose that deficits in the top-down control of endogenous neuromodulation of brain networks result in the deficits of adaptive and flexible cognitive behavior in aging.

## Introduction

The arousal systems of the brainstem, including the noradrenergic, cholinergic and serotonergic systems, shape cognition by altering neural dynamics and promoting synaptic plasticity (Marder, 2012; Froemke, 2015; van den Brink et al., 2019; McCormick et al., 2020; Pfeffer et al., 2021; Jordan, 2024). The function and structure of these brainstem systems changes with aging (Hämmerer et al., 2017, 2018; Dahl et al., 2019, 2022, 2023). While some of these changes have been related to memory deficits (Hämmerer et al., 2017, 2018; Dahl et al., 2019, 2022, 2023), how age-related changes in the brainstem arousal dynamics is related to other aspects of cognition is largely unknown.

Work in younger adults indicates two aspects of cognition that are associated with transient recruitment of brainstem arousal: flexible mappings from sensory input to motor output (Miller and Cohen, 2001; Fusi et al., 2007; van den Brink et al., 2023, 2024) and online inference in changing environments (Nassar et al., 2012; O’Reilly et al., 2013; Filipowicz et al., 2020; Murphy et al., 2021, 2024). In many real-world situations, adaptive behavior requires switching between different stimulus-response associations depending on the context. Context, in turn, often changes in a hidden manner, which then needs to be inferred from ambiguous information (Dayan and Yu, 2006; Filipowicz et al., 2020; Murphy et al., 2021). Adaptive behavior in these situations requires hierarchically nested decisions, in which a higher-order decision process selects the most likely context, which then guides the decision of how to respond in the lower-order process giving rise to overt behavior (Miller and Cohen, 2001; Fusi et al., 2007; Purcell and Kiani, 2016; van den Brink et al., 2024; Calder-Travis et al., 2026).

Several lines of indirect evidence indicate that such hierarchical decision-making may become more challenging with age. First, the ability to perform online inference of hidden changes, based on streams of noisy input data, shows an age-related decline (Nassar et al., 2016; Bruckner et al., 2025). This is in line with the insight that online inference is taxing for working memory and adaptivity (Tavoni et al., 2022), which are in turn sensitive to aging (Bopp and Verhaeghen, 2005). Second, the flexible reconfiguration of stimulus-response associations is taxing for executive control (Miller and Cohen, 2001), which is generally compromised in aging (Cipriani et al., 2025).

The aim of the current study was to identify age-related changes in hierarchical decision-making and its relationship with the dynamic recruitment of brainstem arousal systems. We hypothesized that older adults would show difficulties in online inference of context changes and in the resulting reconfiguration of stimulus response associations. We further predicted that these behavioral difficulties would be associated with an altered task-related recruitment of brainstem arousal systems, specifically in moments of high probability of context switches.

To test these hypotheses, we measured and modeled behavior and arousal dynamics of younger and older adults in a hierarchical decision-making task. Participants had to infer a stimulus-response mapping rule that was undergoing hidden changes (higher-level decision) and apply the inferred rule to report simple visual categorization judgments (lower-level decision). We used pupil dynamics at constant luminance to track the activity of brainstem arousal systems (Murphy et al., 2014; McGinley et al., 2015; Joshi et al., 2016; Reimer et al., 2016; Gee et al., 2017; Breton-Provencher and Sur, 2019; Maheu et al., 2025). Older adults were impaired in the hierarchical decision-making task. Remarkably, the inference of rule switches was largely unaffected, and the age-related impairment mainly reflected the use of incorrect stimulus-response mapping rules. The higher frequency of these occasional rule mistakes in older people was matched by the participants’ subjective experience. While participants’ pupils in both age groups dilated in moments of high rule switch probability, these dilations were smaller in older adults.

## Materials and Methods

### Participants

For the main experiment, data was acquired from 41 younger and 51 older adults. An independent sample of 6 younger and 11 older adults performed an additional control experiment (visual categorization task with fixed stimulus-response mapping rule). One younger participant and three older participants from the main experiment were excluded from the analyses for the following reasons: one younger participant and one older participant presented scores in the Montreal Cognitive Assessment (MoCA), a screening tool for mild cognitive impairment, more than 2 standard deviations (SD) below the mean expected for their age and education level, according to the normative data for the Portuguese population (Freitas et al., 2011); one older participant showed signs of early cataracts and corrected binocular visual acuity less than 8/10; inspection of the anatomical MRI data from one older participant revealed congenital cerebral malformations. The included participants had no history of neurological or psychiatric disease, no history of current alcohol or drug abuse, and were not taking any psychotropic medications. Current medication with beta-blockers was an exclusion criterion. The characteristics of the participants included are described in Table 1.

**Table 1.** Characteristics of participants included in the main experiment (M) and control experiment (C).

|  | N | **Age (years)** |  |  | **Sex** |  | **Education (years)** |  |  | **MoCA scores** |  |  | **Handedness** |  |  |
|---|---|---|---|---|---|---|---|---|---|---|---|---|---|---|---|
|  |  | Mean | Min | Max | F | M | Mean | Min | Max | Mean | Min | Max | Left | Right | Mixed |
| **Young M** | 40 | 24 | 19 | 30 | 25 | 15 | 17 | 12 | 22 | 28 | 25 | 30 | 5 | 34 | 1 |
| **Older M** | 48 | 59 | 51 | 70 | 28 | 20 | 17 | 12 | 28 | 27 | 22 | 30 | 3 | 44 | 1 |
| **Young C** | 6 | 24 | 20 | 29 | 4 | 2 | 16 | 14 | 22 | - | - | - | 1 | 5 | 0 |
| **Older C** | 11 | 64 | 55 | 70 | 6 | 5 | 17 | 11 | 23 | - | - | - | 0 | 11 | 0 |

Older participants included in the main experiment underwent a brief ophthalmologic evaluation to exclude participants with gross eye anomalies that can severely affect their vision. Minimum binocular best corrected visual acuity of 8/10 was necessary for study inclusion. All participants showed no risk of glaucoma presenting normal intraocular pressure. One older participant showed eye conversion difficulties and reported difficulty in 3D vision due to strabismus corrected at young age and was included as the task did not require 3D vision. One older participant reported hearing loss in the left ear and was included as the task did not include auditory stimuli. The MoCA (Freitas et al., 2011), the Beck Depression Inventory II (BDI-II), a depression scale (Beck et al., 1996), and the Wechsler Memory Scale – Third Edition (WMS-III) subtest Logical Memory (episodic memory) (Wechsler, 2008) were used as screening measures. Participants showed no indication of severe depressive symptoms according to the BDI-II. Older participants showed preserved episodic memory with the scores from the Logical Memory test all within the expected for their age, according to the normative data for the Portuguese population, confirming no signs of cognitive impairment (Wechsler, 2008). Three participants (one younger and two older) did not perform the cognitive evaluation due to lack of availability but were still included. The participants of the control experiment met the same inclusion criteria as the participants of the main experiment but did not perform the neuropsychological or the ophthalmological evaluation.

Group sizes differed (41 younger adults and 51 older adults) to compensate for the participants excluded from the behavioral analysis (as described above), and to compensate for participants who were included in the behavioral analysis, but who could not take part in the MRI. We recruited 8 more older participants for the study to compensate for 8 older participants that could not do the MRI session due to difficulties in appropriately correcting vision in the scanner (N = 3), claustrophobia (N = 3), not fitting comfortably in the scanner due to large constitution (N = 1), or lack of availability (N = 1).

The study was approved by the Ethics Committee of the Faculty of Medicine of the University of Coimbra (CE-110/2019). Informed consent was obtained from all human research participants, after explaining verbally and in writing the purpose and possible consequences of the experimental design. The privacy rights of human participants have been observed.

### Procedures

All participants of the main experiment completed at least three experimental sessions. Experimental sessions took place on different days. In session one, participants performed a training block lasting around 30 min where the hierarchical decision-making task was explained step-by-step to ensure that participants clearly understood the task before proceeding (see “Task training procedure” section below). After completing the training, the participants performed four blocks of the hierarchical decision-making task in the psychophysics laboratory while behavioral and eye tracking data were acquired. Participant’s head position was stabilized by using a chin rest to help reduce artifacts caused by head movements. In session two, we acquired structural and fMRI data. The fMRI scans were acquired while participants were engaged in the hierarchical decision-making task with simultaneous eye-tracking acquisition. Four blocks of the task were acquired in the MRI scanner. In session three, participants underwent a neuropsychological evaluation.

At the end of each block, participants received feedback regarding their performance, in the form of the percentage of correct choices in that block written on the screen. In the blocks acquired in the psychophysics laboratory (4 blocks of the hierarchical decision-making task), after receiving the accuracy feedback, participants were also asked verbally if they were aware of responses where, after responding, they immediately realized that they had pressed the wrong button (response lapses). If the answer was “yes”, we subsequently asked them to estimate how many such lapses they experienced. The responses were recorded on paper. In 7% of the total number of task blocks, the responses were not recorded due to time constraints, however, all participants reported their perception of having committed lapses for at least one task block.

For the main study, we combined the behavioral and pupil data acquired in the laboratory with the data acquired in the MRI scanner (4 blocks acquired in the psychophysics laboratory and 4 blocks acquired in the MRI scanner). We observed no significant differences in task accuracy across sessions suggesting stable task performance after the training procedure [repeated measures ANOVA showed no significant effect of acquisition site *F*_(1, 78)_ = 0.008, *p* = .928, or interaction between acquisition site and group *F*_(1, 78)_ = 0.507, *p* = .479]. Analyses of the MRI data will be reported elsewhere.

### Behavioral tasks

#### HIERARCHICAL DECISION-MAKING TASK

The task for the main experiment was a hierarchical decision-making task, which was presented in blocks of 10 min each (Fig. 1). This task required participants to report a visual categorization judgment (lower-order decision), the discrimination between faces and houses, in trials of 2 s duration, with long intertrial intervals (ITIs) ranging from 6.4 s to 22.4 s (see below). These trials are henceforth referred to as “response trials” (Fig. 1). At any moment in between two response trials, the stimulus-response mapping rule required for reporting the lower-order decision could change in a hidden fashion between two possible rules. The rules mapped the two visual stimulus categories (faces and houses) to either left or right button presses (Fig. 1A). To infer which of the two rules was active at any given time, participants were required to monitor a sequence of noisy sensory evidence for one rule over the other (higher-order decision). This evidence was conveyed by the vertical positions of small dots presented in rapid succession every 400 ms along the vertical meridian (Fig. 1B; duration of each evidence sample: 100 ms). The vertical positions of the dots were drawn from one of two generative distributions, which were in turn associated with one of the two stimulus-response mapping rules (Fig. 1C). The two generative distributions were two Gaussians with equal standard deviation (σ_upper_ = σ_lower_ = 0.4° visual angle) and different means symmetric around fixation (|μ_upper_| = |μ_lower_| = 0.25° visual angle from the center of the screen). Each dot position provided evidence of which generative distribution was active (log likelihood ratio for one generative distribution over the other) and, therefore, will be referred to as an evidence sample. The generative distribution could change from one evidence sample to the next with a low probability (hazard rate) of 1/70. The process generating the evidence samples was reset at the end of each block. Response trials were inserted between evidence samples if either of these conditions was met: (i) a successful coin flip based on a probability of success of 1/35 and at least 11 new evidence samples presented since the last response trial or (ii) the number of evidence samples since the last response trial exceeded 50. Thus, between response trials there were a minimum of 11 and maximum of 51 evidence samples and the ITIs varied between 6.4 and 22.4 s (including the 2 s duration of the response trial). In response trials, images of faces or houses (the stimuli for the lower-order decision) were presented for 0.5 s followed by 1.5 s fixation-only screen. Participants were instructed to report the image’s category within the 2 s response window from image onset with their left or right index fingers with the ‘z’ or ‘m’ keys from the computer keyboard in the blocks acquired in the laboratory and with MRI-compatible button interfaces positioned on each side of the body in the blocks acquired in the MRI scanner. Response trials with no response or with reaction time longer than 2 s were considered missed trials. The accuracy of the response depended on the selection of the correct rule and the correct identification of the stimulus category. It is important to note that the stimuli were clearly visible, and the participants did not report any difficulty discriminating between faces and houses as confirmed in the control task (see section “Visual categorization task with fixed stimulus-response mapping rule” and Fig. S1). Thus, accuracy in the hierarchical decision-making task depended mostly on the selection and implementation of the correct stimulus-response mapping rule and not on the identification of the visual category of the target stimuli. The relationship between the two generative distributions and the mapping rules (counterbalanced across participants) stayed constant across all experimental sessions (Fig. 1C; younger: sub-group 1 N = 20, sub-group 2 N = 20; older: sub-group 1 N = 23, sub-group 2 N = 25). Participants were reminded of which distribution corresponded to which rule at the beginning of each block with a visual representation of the stimulus-response mapping rules.

**Figure 1.**
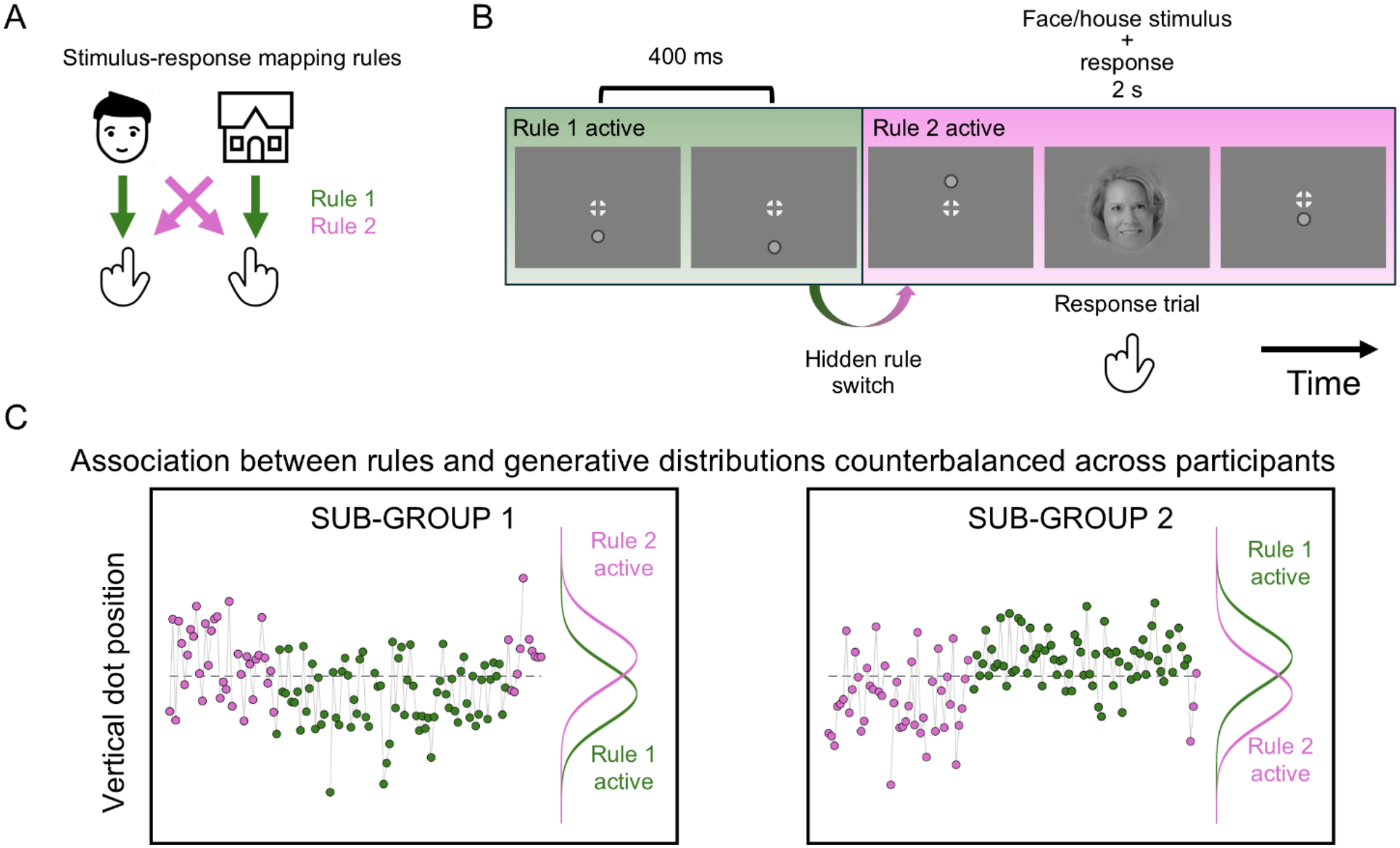
Hierarchical decision-making task. **A.** Schematic of task stimulus-response mapping rules. In the visual categorization response trials, participants were required to respond to images of faces or houses according to one of two stimulus-response mapping rules, Rule 1 (green) or Rule 2 (pink). The images (represented in **A** as line drawings) were grayscale photographs of faces or houses on grey phase-scrambled backgrounds presented on a circular aperture (Methods). **B**. Schematic (not to scale) of the sequence of evidence samples (stream of small dots presented one-by-one around the fixation target along the vertical meridian) preceding and following a response trial. Each dot was an evidence sample of which stimulus-response mapping rule was active. Dots were presented for 100 ms and the time from one dot onset to the next was 400 ms. Response trials consisted of images of faces or houses presented for 500 ms, followed by a blank screen with the fixation target only for 1500 ms before the next evidence sample. **C**. Evidence samples were drawn from one of two generative distributions (Gaussians with equal standard deviation and means symmetric around fixation) producing a noisy evidence stream. The active generative distribution governed the active rule and could switch unpredictably between any two evidence samples (probability: 1/70). The association between the generative distribution and the active rule was counterbalanced across participants with approximately half of the participants associating the lower distribution to Rule 1 and the upper distribution to Rule 2 (sub-group 1), and the other half associating the upper distribution with Rule 1 and the lower distribution to Rule 2 (sub-group 2).

The task was created using MATLAB (The MathWorks Inc.) and the Psychophysics Toolbox Version 3 (Brainard, 1997) and was presented on a grey background. A fixation target was presented at the center of the screen throughout the task, except when the images of faces or houses were briefly displayed. The fixation target consisted of a white combination of bull’s eye and crosshair shape located in the center of the annulus, as recommended by Thaler et al. (2013) for experiments that require stable fixation (Thaler et al., 2013).

In the laboratory, the stimuli were presented on a computer monitor (19-inch Dell monitor) with a spatial resolution of 1440 x 1080 pixels, a refresh rate of 100 Hz, and 52.5 cm in width and 39.5 cm in height. In the MRI-scanner, stimuli were presented on an MRI-compatible LCD-screen, which the participants viewed through a mirror mounted above their eyes, with a spatial resolution of 1920 × 1080 pixel, a refresh rate of 60 Hz, and 87.8 cm in width and 48.5 cm in height. Correction to normal vision in the scanner was ensured using magnetic field-compatible eyeglasses.

The stimuli used in the visual categorization trials were face and house stimuli included in the fLoc functional localizer package (Stigliani et al., 2015). From their stimulus sets, we selected images of faces and houses that were clear representatives of that category. To ensure that the categories could not be identified based on luminance differences, we measured the luminance of the center of each image using a spectroradiometer (Apacer Technology Inc). We excluded images with luminance with absolute *z*-score higher than 1.5. The luminance values of the final image sets, comprising 64 face and 65 house images, were not significantly different across categories. The images were displayed through a circular aperture in the center of the grey screen.

#### VISUAL CATEGORIZATION TASK WITH FIXED STIMULUS-RESPONSE MAPPING RULE

A control experiment was conducted in a separate cohort of participants (Table 1) to verify that (as intended) the images of faces and houses used in the main task were easy to discriminate and categorize for both younger and older adults. These participants performed two blocks of a visual categorization task (Fig. S1A). In each block, we presented in random order all the face and house stimuli used in the hierarchical decision-making task (64 face and 65 house images) (Stigliani et al., 2015). The participants were instructed to report if they perceived a face or a house by pressing a key on a computer keyboard with their left or right index fingers within a 2 s time window starting at stimulus onset. The stimulus-response mapping rule was fixed for each participant and counterbalanced across participants (approximately half of the participants responded with left index finger for faces and right index finger for houses and *vice versa*). As in the hierarchical decision-making task, the images were presented at the center of the screen on a circular aperture for 0.5 s followed by a grey screen presented for 1.5 s. The ITI was between 1 s and 2.5 s randomly drawn independently from a uniform distribution. A central fixation target consisting of a white combination of bull’s eye and crosshair shape located in the center of the annulus (Thaler et al., 2013) was presented at all times except when the images were presented. Participants performed a short training version of the task (20 trials) to ensure they understood the instructions, followed by two task blocks, with a self-paced break in between blocks.

#### TASK TRAINING SCHEME INCLUDING INFERENCE-ONLY, INSTRUCTED RULE AND HIERARCHICAL TASK

A step-by-step training scheme comprised of 5 phases was used to ensure the participants understood the requirements for the hierarchical decision-making task. During training, response trials were followed by written feedback (the expressions “correct”, “wrong”, or “you did not respond” written in Portuguese) displayed on the screen 2 s after stimulus onset. Feedback stayed on the screen until a key was pressed to continue the task. Participants were encouraged to report any difficulties and ask any questions. Each training phase included on average 12 response trials (400 evidence samples), except phase 1 that was very easy (as described below) and only included an average of 2.5 response trials (100 evidence samples). Task instructions and the training phase were repeated if difficulties were recognized. In phases 1, 2 and 3, participants performed a version of the task that tested the ability to infer which generative distribution was active, the inference-only task. This task had the same characteristics as the hierarchical decision-making task, with the difference that the response trials were signaled by the presentation of a grey circle surrounding the fixation target instead of the images of faces and houses (Fig. 2F). The grey circle was presented for 0.5 s followed by only the fixation target for 1.5 s. The response window was 2 s from stimulus onset. Participants were instructed to report on response trials which generative distribution they believed was active at that moment using their right index and right middle fingers and the keyboard keys ‘left arrow’ (for lower distribution) and ‘up arrow’ (for upper distribution). Phases 1 and 2 used an easier version of the inference-only task with lower overlap between the generative distributions (standard deviation of 0.4° visual angle and means located at 0.5° visual angle above and below the center of the screen along the vertical meridian). In phase 1, evidence samples were additionally colored according to which generative distribution was generating them (green for upper distribution and blue for lower distribution). In phase 2, participants performed the inference-only task without the colored evidence samples. In phase 3, the participants performed the inference-only task with the final parameters of the generative distributions (standard deviation of 0.4° visual angle and means located at 0.25° visual angle above and below the center of the screen along the vertical meridian). In phase 4, we trained the application of the two stimulus-response mapping rules (Fig. 1A) by using a version of the hierarchical decision-making task with colored evidence samples making explicit which generative distribution was active at any time by the dots color (task with instructed rule: Fig. 2F). Participants now responded to the face and house stimuli in response trials with their left and right index fingers using the ‘z’ and ‘m’ keys from the computer keyboard. In phase 5, we trained the full version of the task (named inferred rule in Fig. 2F and G). The rationale for having different response rules for the different tasks during training (right index and middle fingers for the inference-only task and right and left index fingers for the full task) was to avoid creating interference between the representation of the (upper or lower) sources of the evidence samples and the representation of the rule (i.e. mapping between left vs. right hand actions and faces vs. house stimuli). Training data for 1 older and 1 younger participant and the data from the first 3 phases of training for 1 younger participant were lost due to recording errors.

**Figure 2.**
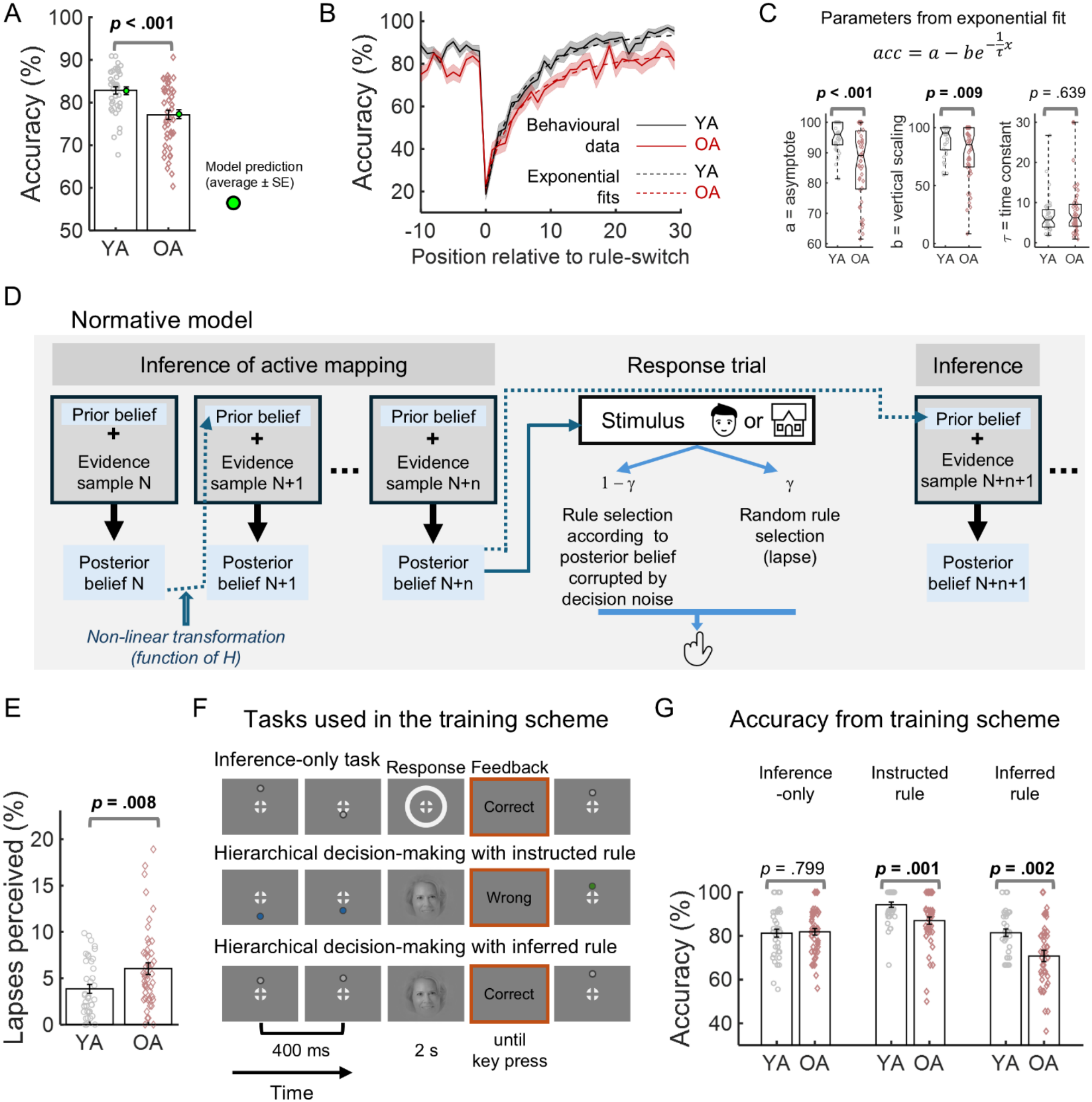
Age effects on task performance. **A**. Accuracy of younger (YA) and older adults (OA), and the best-fitting model (green circles, see Methods). **B**. Accuracy locked to hidden rule switches as a function of the number of evidence samples presented since the rule switch. Continuous black and red lines, average for YA and OA; shading, standard error of the mean (SEM). Dashed black and red lines, average of exponential fits to the YA and OA data. **C**. Parameter estimates from exponential fit. *P*-values from Mann–Whitney U test. On the boxplots, the central mark indicates the median, and the bottom and top edges of the box indicate the 25th and 75th percentiles, respectively. The whiskers extend to the most extreme data points. **D.** Schematic of the normative model. **E.** Lapses reported by the participants displayed as percentage of total number of response trials (see main text). **F.** Schematic (not to scale) of the tasks used in the training scheme designed to ensure that the participants understood the hierarchical decision-making task. Each training task included on average 12 response trials. **G.** Participants’ performance in the training tasks. Data points, individual participants; bars, group average; error bars ± SEM. *p*-values in A, E, and G from independent samples *t*-tests.

### Behavioral analyses

For task accuracy and reaction time calculation, we did not consider the visual categorization trials where participants responded with more than one key press or did not respond within 2 s from stimulus onset. Eight older participants did not perform the fMRI acquisition, as explained above, and older participants were more likely to respond with more than one key press invalidating the trial. For these reasons, the older group had on average lower number of valid response trials (mean ± SD: younger = 292 ± 32; older = 259 ± 59; *t*_(86)_ = 3.15, *p* = .002).

To quantify the effect of rule-switches on task accuracy, we calculated the average accuracy as a function of the distance of the visual categorization trial from a rule-switch (measured in number of evidence samples presented). Missing accuracy values were replaced by the average value from the corresponding age group for that trial position. To quantify age group differences, we fitted the exponential function

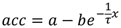

to each participant’s accuracy data as a function of *x*, the evidence sample from rule-switch. The free parameters of the exponential function were the asymptote *a,* the vertical scaling *b*, and the time constant *τ*. Curve fitting was performed for each participant data separately in MATLAB, and the free parameters were constrained within the upper bounds of 100 for asymptote, 100 for vertical scaling and 30 for the time constant and lower bounds of zero for all parameters.

### Behavioral modeling

We fit a family of computational models to the behavioral data from the hierarchical decision-making task (Calder-Travis et al., 2026) using code written in MATLAB.

#### NORMATIVE MODEL

In the hierarchical decision-making task, participants must combine the noisy evidence samples in order to determine which of two states is currently active, corresponding to the two possible generative means and thereby also corresponding to the active stimulus-response mapping rule. Hidden changes in state can occur between the evidence samples with a probability determined by a fixed hazard rate (H). The optimal solution to this problem is known (Glaze et al., 2015) and involves, at each new evidence sample, updating the prior about the current state using the log-likelihood ratio corresponding to the new evidence sample. This gives the log-posterior ratio, which is non-linearly transformed into the log-prior ratio for the next evidence sample (reflecting the fact that a hidden change in state may occur before the next evidence sample is presented).

In the equation (Glaze et al., 2015):

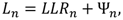

where *L_n_* indicates the log-posterior ratio for one state over the other after observing evidence sample *n*, *LLR_n_* indicates the log-likelihood ratio corresponding to evidence sample *n*, and **Ψ***_n_* indicates the log-prior ratio, which can be computed through,

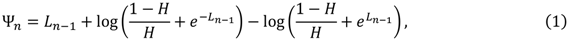

where *H* indicates the hazard rate.

Denote the position of a dot, *x_n_*, where *n* indicates that this is the *n*th evidence sample in the block, and the position along the vertical meridian is measured in degrees of visual angle from the centre of the screen. Recall that dots positions follow a normal distribution, with the mean of the normal distribution depending on the active state. Using this information, the log-likelihood ratio can be computed through (Glaze et al., 2015),

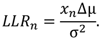

Here Δ*μ* indicates the difference between the generative means of the dots’ vertical position under the two different states, and *σ* corresponds to the standard deviation of the normal distribution that the dots’ vertical positions are drawn from.

The participant’s belief is a form of decision variable in that it tracks the developing evidence and prescribes which mapping rule to use or which response to make based on whether the decision variable is positive or negative (Gold and Shadlen, 2001). We allow the possibility that the mechanism that translates the participant’s decision variable (*i.e*., belief) to a mapping might be noisy, and we consider two sources of noise. First, we allow the possibility that during the readout of the decision variable, the readout is corrupted by normally distributed noise. Denote the stimulus-response mapping used in the hierarchical task, R, and the two mappings, R=1 and R=0. In the absence of any other further sources of noise beyond normally distributed noise, the probability of the participant using mapping R=1 would be:

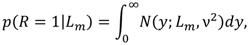

where ν is the standard deviation of this “readout” noise, and *m* is the index of the most recently presented evidence sample.

However, we additionally model the possibility that with some specific probability, *γ*, a “lapse” is made, resulting in a random mapping being selected. This gives us the following final equation for the probability of the participant using mapping R=1:

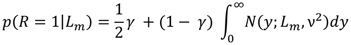

This model has two free parameters, the standard deviation of the normally distributed noise, ν, and the probability of a lapse, *γ*. In the main text, we refer to the above model as the “normative” model.

#### ALTERNATIVE MODELS

We considered a number of alternative models. All model variants featured the two sources of noise from the normative model described above, and hence all feature the two associated free parameters. The models differ in how the decision variable is computed.

We used a variant of the normative model, “normative free H”. In this model the computation of the decision variable is almost unchanged, but we relax the assumption that participants have learned the true hazard rate (Murphy et al., 2021). The equations for this model are the same as the normative model, except that in the equation for computing the log-prior ratio (equation 1) participants use their subjective estimate of the hazard rate, *H*^, rather than the true value. For this model, we fit subjective hazard rate as an additional free parameter (3 free parameters in total).

We considered a number of other non-normative models, that represent simpler heuristic strategies (Murphy et al., 2021). In the “last sample” model, the decision variable is simply the value of the most recent evidence sample, *x_m_*. All other samples are disregarded (2 free parameters in total). In the “perfect accumulator” model, the observer sums all evidence samples since the start of the current block of the task and uses this total value as their decision variable (2 free parameters in total).

The “bound accumulator” is a variant of the “perfect accumulator”. The observer again sums the value associated with every new evidence sample that they receive (since the start of the current block of the task), but there is now an upper and lower bound on the value (Θ and −Θ) that the accumulator can take (Glaze et al., 2015). The accumulator cannot go above or below these bounds, meaning that if the accumulator reaches the upper bound, Θ, it remains at Θ with further positive having no further effect, until negative evidence pushes the state of the accumulator below this bound. The accumulator is again used as the decision variable (3 free parameters in total).

The “leaky accumulator” is a further variant of the “perfect accumulator” (Glaze et al., 2015). Each time a new evidence sample is presented, before the new evidence value is added to the accumulator, some proportion of the information in the accumulator is discarded. Specifically, the state of the accumulator is multiplied by, (1-α), where α is a number between 0 (no leak) and 1 (total leak). This gives 3 free parameters in total.

#### MODEL FITTING

Models were fit on a participant-by-participant basis using maximum likelihood fitting (specifically, negative log-likelihood was minimised using the MATLAB “fmincon” algorithm (Matlab optimization toolbox (Version 8.0), MathWorks, 2017). Fitting was repeated 10 times for each participant and model. We only retained and used the best fit of these 10 fits (again evaluated using the likelihood).

Each fit began by evaluating the likelihood of 250 candidate starting points. Each candidate start was constructed by drawing values for all relevant parameters from uniform distributions. The uniform distribution for a parameter ran from the “Lower initial” value to the “Upper initial” value specified in Table 2. The value for “Upper initial” for ν was 6, except for the perfect accumulator model where we increased this to 200, because the decision variable has a tendency to reach large values.

**Table 2.** Upper and lower bounds used during fitting and for initially drawing candidate start points.

| Parameter | Lower bound | Lower initial | Upper initial | Upper bound |
|---|---|---|---|---|
| *Ĥ* | 0 | 0.001 | 0.5 | 1 |
| ν | 0 | 0.005 | 6 or 200 | 2000 |
| *γ* | 0 | 0 | 0.4 | 1 |
| Θ | 0 | 0.1 | 12 | 2000 |
| α | 0 | 0 | 1 | 1 |

The candidate start point with the greatest likelihood was used as the starting point in the “fmincon” algorithm. During fitting, free parameters were constrained to be within the “Lower bound” and “Upper bound” values shown in Table 2.

We did not use the training data for fitting the models, nor did we use trials with no response within the 2 s response window (missed trials) or trials with more than one button press within the response window (invalid trials). Fitting was conducted using all hierarchical decision-making task data from both the psychophysics lab and the scanner.

#### MODEL COMPARISON AND DATA SIMULATION

We calculated the Akaike information criterion (AIC) and Bayesian information criterion (BIC) separately for each participant and model. The normative free H model (the model with lower AIC and BIC when considering all participants) was used as reference for comparison and ΔAIC and ΔBIC were calculated as differences from the reference model. ΔAIC and ΔBIC were considered meaningful if these values were significantly different from zero assessed with one sample *t*-tests (Fig. S2).

Additionally, we simulated data from the models in order to see how well they capture features of the real data. We performed simulations on a participant-by-participant basis, simulating the same number of trials as the participant actually performed, and using the stimuli that were actually presented to that participant. Using a specific model, the fitted parameters for a participant, and the stimuli shown, we could estimate the time-course of the decision variable. In turn, this allowed us to compute the probability, according to the model, that the participant would use each rule (hierarchical decision-making task). Finally, we draw rules according to these probabilities and calculated the model’s accuracy.

#### MODEL-BASED LATENT VARIABLE ESTIMATION

Using the normative model (with the hazard rate set to the true value of 1/70) we computed, at each evidence sample, two computational variables of interest.

We computed a measure of uncertainty using the negative of the absolute value of the log-prior ratio (Murphy et al., 2021): −|**Ψ***_n_*|. We also computed the probability that the state had changed between the previous evidence sample, n-1, and the current evidence sample, n (change point probability, CPP). This quantity can be computed as follows (Murphy et al., 2021):

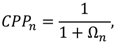

where,

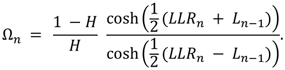

### Pupil data acquisition and preprocessing

The EyeLink 1000 Plus desktop mount (SR Research, Ottawa, Ontario, Canada) was used to record the horizontal and vertical gaze position and the pupilogram. A trigger pulse was generated at each stimulus onset and button-press. The data was recorded monocularly at a sampling rate of 500 Hz. Eye position calibration and validation were conducted before each block. Eye-tracking inside the MRI scanner was technically more challenging. For a small number of participants not all blocks acquired in the scanner had usable eye tracking data due to difficulty of the eye tracker detecting the eye/pupil through glasses: 1 younger and 1 older (from the 40 that did the MRI acquisition session) did not have any usable eye tracking data in the scanner; 1 younger participant only had 1 run with usable eye tracking data in the scanner; and 1 older participant only had 3 runs with usable eye tracking data in the scanner.

Pupil data was analyzed with custom MATLAB scripts and the EEGLAB toolbox (Delorme & Makeig, 2004) in MATLAB (The MathWorks Inc. MATLAB Version: R2024b, Natick, Massachusetts). Code used for pupil preprocessing was adapted from (Urai et al., 2017). Missing data and blinks, as detected by the EyeLink software, and additional blinks identified using peak detection on the velocity of the pupil signal, were padded by 150 ms and linearly interpolated (Urai et al., 2017). Pupil data were normalized within each block to the percentage of the mean, band-pass filtered between 0.06 and 6 Hz, and down sampled to 50 Hz.

### Task-related evoked pupil responses

We studied the pupil responses locked with response trials, rule switches, periods of stable rule (no rule switch), and high or low CPP. To isolate the signals of interest and correct for the temporal overlap between the responses to the different events in the continuous pupil data, we used linear-regression-based deconvolution modeling. Deconvolution allows for the correction for overlap between temporally adjacent responses (Dale, 1999).

For the study of the effect of rule switch on the pupil, we modeled three events: rule switch (the onset of the evidence sample where the rule switch occurred), no rule switch (defined as the evidence sample halfway between two rule switches associated with a period where the rule was stable), and the onset of the response trial (the onset of the face or house image). This allowed us to compare the pupil responses to evidence samples associated with a rule switch versus samples associated with a stable rule period while controlling for the pupil response to response trials. Each event was modelled from 5 s before up to 15 s after.

For the study of the effect of CPP on the pupil, we modeled three events: high CPP, low CPP, and response trial (the onset of the face or house image). Evidence samples with CPP higher than 0.0775 were considered high CPP events and samples with CPP lower than 0.000825 were considered low CPP events. These CPP thresholds were calculated so that the across participants average number of high CPP and low CPP samples were similar to the number of rule switches. Average number of rule switches (SD) = 215 (65); average number of high CPP events (SD) = 214 (62); average number of low CPP events (SD) = 216 (65).

For each analysis, we deconvolved the mean responses (Dale, 1999; Donner et al., 2008) locked to the three event types according to:

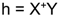

where Y was the pupil time series, h = [h_1_*^T^* h_1_*^T^* h_1_*^T^*]*^T^*was a vertical concatenation of estimated responses locked to the three event types with the superscript *T* denoting matrix transpose, the design matrix X = [X_1_ X_2_ X_3_] was a horizontal concatenation of three convolution matrices corresponding to the three event types, and + denotes the pseudoinverse. Each X*i* had dimensions *M* x *N*, where *M* was the length of the pupil time series and *N* was the number of time points in the estimated h*i* from 5 s before the event up to 15 s after. The first column of each X*i* contained 1’s at the samples corresponding to 5 s before the event and 0’s elsewhere. Each subsequent columns contained 1 in the position corresponding to the following time point and 0’s elsewhere forming a matrix with a diagonal of 1’s corresponding to the time points from 5 s up to 15 s after the event.

Statistical analyses of the deconvolved pupil responses were performed time point-by-time point using the LIMO EEG toolbox, an open-source MATLAB toolbox (Pernet et al., 2011). We ran second level analysis (group level) using robust trimmed means methods. Results were reported corrected for multiple testing using a temporal clustering forming threshold of *p* = 0.05 (Pernet et al., 2015). Cluster *p*-values were calculated by determining the maximum cluster-mass (sum of the statistical values within a cluster) of each of 1000 bootstraps (in which each condition/subject is first mean centered, then sampled with replacement) and calculating the proportion of bootstraps that resulted in a larger cluster-mass than the cluster-mass of the observed cluster (Maris and Oostenveld, 2007; Pernet et al., 2015). To compare the deconvolved pupil responses locked with response trials across groups, we used independent samples *t*-tests comparing the responses calculated in the deconvolution. The specific deconvolution used featured the effect of response trials modelled together with the effect of rule switches including the predictors for the rule switches and the predictors between the rule switches (no rule switch). To compare the effects of rule switch or CPP across age groups, the deconvolved responses were analyzed using repeated measures ANOVAs with group as between-subjects factor and rule switch/no rule switch or high CPP/low CPP as within-subject factor. To investigate the association between task accuracy and the pupil responses to CPP or pupil responses in response trials, we ran time point-by-time point across subjects’ regression analyses investigating the association between accuracy and the amplitude difference between the pupil responses locked with high CPP and low CPP samples or the amplitude of the response in response trials locked with stimulus onset. The regressions were run separately for each age group.

To compare the pupil responses across acquisition site (laboratory *versus* MRI scanner), we run the deconvolution separately for the blocks acquired in the laboratory or in the scanner and run time point-by-time point repeated measures ANOVA as described above but including acquisition site as an extra within-subject factor. For these analyses, we only included participants that had pupil data in the laboratory and in the scanner (N = 39 young and N= 39 older). Similarly, to compare the pupil responses in the first half of the task (blocks 1 and 2 from each session) with the responses in the second half of the task (blocks 3 and 4), we run the deconvolution analyses separately and run repeated measures ANOVA including time-on-task (blocks 1 and 2 *versus* blocks 3 and 4) as an extra within-subject factor.

### Assessment of general cognitive function

General cognitive function was assessed with a set of 9 standardized neuropsychological tests in a session of approximately 1.5 h duration. The (MoCA) (Freitas et al., 2011), the BDI-II, and the WMS-III subtest Logical Memory (Wechsler, 2008) were used as screening measures. The remaining 6 tests consisted of 3 sub-tests of the WMS-III (Wechsler, 1987, 2008) and 3 tests from the CANTAB Connect Research (Cambridge Cognition Ltd) run on an iPad (Supplementary Table 1). The tests were selected to assess four cognitive domains: processing speed, working memory, resistance to interference, and planning. Processing speed was assessed with the CANTAB Reaction Time (RTI) test. Working memory was assessed with the WMS-III, subtests Spatial Span, Digit Span and Letter Number Sequence. Resistance to interference was assessed with the CANTAB Multitasking (MTT) test. Planning was assessed with the CANTAB One Touch Stockings of Cambridge (OTS) test. In the older group, the data for the Logical Memory test was lost for one participant due to difficulty recovering recorded material and for another participant the responses during the CANTAB MTT test were not correctly registered due to technical issues. We performed an empirical grouping by conducting a principal-components factor analysis with varimax rotation and Kaiser normalization, on the centered test measures, using IBM SPSS Statistics (version 27). We grouped the test measures with rotated factor loadings of .50 or higher on the same composite (Supplementary Table 2) (Wilson et al., 2003). Measures that loaded on more than one factor were assigned according to the highest loading. Measures without loadings of .50 or higher were not included. Factor analyses revealed that the measures from the cognitive tests were best grouped in three composites that we interpreted as Working Memory Composite (including the measures from the three working memory tests), Processing Speed Composite (including the measures from the RTI test and the latency measure from the MTT test), and Executive Control Composite (grouping measures from the OTS and MTT tests). To compute the composite scores for each cognitive domain, raw test scores were converted to z-scores, using the mean and standard deviation from all participants, and the z-scores were averaged. The composite scores were calculated so that higher scores were associated with better cognitive function. The z-scores of the measures where higher scores meant worse performance were multiplied by –1 before averaging.

To investigate if variability in executive control explained the task accuracy difference across age groups, we run a regression including all participants with accuracy as independent variable and executive control composite score as predictor to calculate the accuracy residuals after controlling for the difference in executive control. We then compared the residuals across age groups using independent samples *t*-tests.

### Within-analysis control for multiple comparisons

For the analyses involving several parameters or tasks, we used Bonferroni correction to adjust for multiple comparisons by multiplying the resulting *p*-values by the number of parameters or tasks tested (adjusted *p*-value) and assuming an alpha level of .05.

### Code Accessibility

Custom code used in the study is available on Github: https://github.com/mariajribeiro/sensory_motor_reconfiguration_in_aging

## Results

We measured behavior and pupil dynamics in a group of older adults (N = 48) and a group of younger adults (N = 40) performing a hierarchical decision-making task (Figure 1). The lower-level decision required participants to categorize photographs as faces or houses and report their categorization judgment by pressing one of two buttons. The higher-level decision was the internal selection of the stimulus-response mapping rule used for reporting the lower-level decision (Fig. 1A). The active rule could undergo hidden changes at any moment with a low probability (hazard rate). To infer the active rule, participants tracked a rapid sequence of “sensory evidence samples” (Glaze et al., 2015; Murphy et al., 2021), which was presented in the format of visual dots flashed along the vertical meridian, during the long and variable (6.4 to 22.4 s) intervals between the response trials of the lower-level decision. The dot positions were drawn from one of two overlapping Gaussians with equal standard deviations and a mean symmetrical about the horizontal meridian (see Methods). Each distribution was associated with one stimulus-response mapping rule (Fig. 1C), so that each dot provided a sample of evidence for one over the other rule being active (Glaze et al., 2015; Murphy et al., 2021).

### Impaired hierarchical decision-making in older participants

While both groups of participants were able to perform the hierarchical decision-making task above chance level, the older group had lower accuracy than the younger group (mean ± SD: younger = 83 ± 5 %; older = 77 ± 7 %; *t*_(86)_ = 4.12, *p* < .001; Fig. 2A). The older participants were also more likely to respond with more than one keypress, possibly reflecting correction of premature choices [median (interquartile range, IQR) percentage of response trials with more than one keypress: younger = 0.3% (0.7%), older = 0.7% (2.2%); Wilcoxon Rank Sum Test *p* = .002]. We found no evidence for a difference between groups in the percentage missed trials [median (IQR): younger = 0% (0.4%), older = 1% (1.2%); Wilcoxon Rank Sum Test *p* = .142] or in reaction time [mean ± SD: younger = 816 ± 15 ms; older = 830 ± 11 ms; *t_(_*_86)_ = –0.509, *p* = .612]. A control experiment indicated that the face and house stimuli were well discriminable for both age groups, yielding near-ceiling performance (Fig. S1). This indicates that the image categorization was neither a performance-limiting factor in general nor a source of any age group differences in the hierarchical decision task.

To shed light onto the sources of errors underlying the age-related performance difference, we quantified the temporal dynamics of accuracy following rule switches. As expected, the latter was followed by a marked dip in performance, which then recovered over time as more evidence in favor of the new rule was accumulated (Fig. 2B). One possibility is that older adults were slower in this recovery, indicating that they were impaired in detecting the hidden rule changes, in implementing the resulting behavioral change, or in a combination of both. An alternative possibility is that older participants were impaired in their steady-state performance independently of the rule switches. To arbitrate between these alternatives, we fitted an exponential model to the accuracy data following the rule switch. Both groups recovered their accuracy at a similar rate [time constant *τ* was not significantly different across groups – median (IQR): younger = 6.8 ± 4.6; older = 10.4 ± 21.6; Mann-Whitney U test *z* = –0.469, *p* = .639]. By contrast, the older group’s accuracy converged to a lower value reflected in a significantly lower asymptote (*a*) and vertical scaling parameter (*b*) [median (IQR) asymptote: younger = 96 (7); older = 89 (19); Mann-Whitney U test *z* = –3.84, *p* < .001; vertical scaling: younger = 95 (19); older = 85 (34); *z* = –2.60, *p* = .009; Bonferroni adjusted *p*-values asymptote *p_adjusted_* < .001 and vertical scaling *p_adjusted_* = .027, see Methods].

Because our control results excluded the lower-order decision in the context of fixed stimulus-response mappings as a relevant source of errors, the lower asymptotic performance of older participants points to a more frequent use of the incorrect rule in the context of the hierarchical decision-making task. In sum, older adults could categorize the images and also detect, and react to, rule changes as well as younger adults, but they seemed to be impaired in implementing the required stimulus-response mapping rule in the context of a task that required rule switches. This interim conclusion is further supported by both, computational modeling of their behavior as well as participants subjective reports, as presented next.

### Computational model-inferred and subjectively experienced lapses

Participants’ behavior was well fitted by a computational model, in which the higher-level decision approximated the normative belief updating in volatile environments (Glaze et al., 2015; Murphy et al., 2021) (Fig. 2A, green dots and Fig. S2). In the normative model, each evidence sample (log likelihood ratio, LLR) was combined with a prior belief (Ψ) into a posterior belief (L) for one over the other rule (Fig. 2D). Our model entailed three sources of suboptimality as free parameters: the subjective hazard rate (H) captured biases in the representation of environmental volatility, decision noise corrupted the transformation from the posterior belief to the rule selection, and the lapse rate was the probability of random responses that were independent of belief (i.e. independent of accumulated evidence). Indeed, this model, referred to as “normative free H” model, fitted the data better than a range of alternative models, including heuristic strategies, with lowest mean AIC and BIC (205 and 216, respectively; both 0.51 smaller than next-lowest value, the leaky accumulator model; see Figure S2A-B for model comparisons split by group).

The estimated model parameters pointed to a higher lapse rate, but not decision noise, in the older than the younger group (Fig. S2C). The group effect on lapse rate was significant for uncorrected *p*-values and marginally significant after correcting for multiple comparisons [lapse rate median (IQR): younger = 0.045 (0.093), older = 0.087 (0.194); Mann-Whitney U test, *z* = –2.18, *p* = .029; BF_10_ = 1.32; *p_adjusted_* = .087; decision noise median (IQR): younger = 1.99 (2.18), older = 3.03 (3.64); Mann-Whitney U test, *z* = –1.89, *p* = .058; BF_10_ = 1.33; *p_adjusted_* = .174]. We found no evidence for significant group differences in H [median (IQR): younger = 0.009 (0.019), older = 0.004 (0.025); Mann-Whitney U test, *z* = –1.00, *p* = .319; BF_10_ = 0.18].

After each block of the hierarchical decision-making task (excluding training blocks and the blocks acquired in the scanner), we asked participants if they were aware of any errors in the preceding block, where they had responded and immediately realized they made an error. If the response was “yes”, we subsequently asked them to estimate how many such lapses they experienced (Methods). Older participants reported a higher fraction of lapses [mean ± SD of lapses as percentage of number of response trials: younger 3.9 ± 3.0 %, older 6.0 ± 4.3 %; *t*_(86)_ = –2.7, *p* = .008; Fig. 2E]. This increase in perceived lapses in older people extends to a significant increase in the ratio of reported lapses to accuracy (i.e. the number of lapses reported by a participant divided by their accuracy) [mean ± SD percentage of lapses per accuracy: young = 5.0 ± 4.1 %, older 8.6 ± 7.5 %; *t*_(86)_ = –2.8, *p* = .007]. In other words, older participants had a stronger tendency than younger participants to perceive that they made errors by pressing the wrong button while knowing the correct decision. The model-based parameter estimates combined with the subjective reports support the conclusion that difficulties in the sustained application of the active stimulus-response mapping rule were a dominant factor governing the reduced accuracy of the older group in the hierarchical decision-making task.

### Specific impairment of older adults in the application of changeable stimulus-response mapping rules

To distinguish between the ability to perform the online inference and the flexibility of switching between different stimulus-response associations (implementation of the volatile stimulus-response mapping rules), we took advantage of the fact that the training scheme used to ensure the participants understood the hierarchical decision-making task included two component versions of the task: an “inference-only” task, in which they explicitly reported the switches in the hidden state without applying it to a rule switch, and an image categorization task with same frequency of rule switches, which were now explicitly instructed (“instructed rule task”; Fig. 2F, see Methods for details). The training ended with a version of the full hierarchical decision-making task, which is referred to as “inferred rule” for comparison in Figure 2F and G. Each training phase had on average 12 trials and was repeated if deemed necessary. We used the data from the last training repetition of each task to compare the performance across age groups. Compared to the younger group, the older group was not impaired in the inference-only task [*t*_(83)_ = –0.256, *p* = .799], but was significantly impaired in both the instructed rule [*t*_(84)_ = 3.38, *p* = .001] and the inferred rule task [*t*_(84)_ = 3.27, *p* = .002]. These comparisons remained significant after adjusting for multiple comparisons taking into consideration the 3 tasks (instructed rule task *p_adjusted_* = .003, inferred rule task *p_adjusted_* = .006). These results are consistent with the idea that the older adults were specifically impaired in the implementation of volatile stimulus-response mapping rules and not in the inference of which mapping was active and the respective change points. Further in line with this idea, older adults also needed to repeat the instructed rule task more often than younger adults during training [median (IQR): young = 1 (1), older = 3 (2); Wilcoxon Rank Sum Test *p* < .001]. The number of training repetitions for the inference-only and inferred rule tasks did not differ across groups [median (IQR): inference-only and inferred rule: 1 (0) for both groups, Wilcoxon Rank Sum Test *p* = .176 and *p* = .080, respectively].

Taken together, behavioral analysis and modeling established an impairment of older people in the hierarchical decision-making task and several lines of convergent evidence support the idea that this is due to a difficulty in implementing in an effective and sustained manner stimulus-response mapping rules in a context that required frequent rule switching (see Discussion). We next tested our hypothesis, that this impairment may be related to an insufficient recruitment of arousal systems by the rule switches in our task.

### Altered recruitment of pupil-linked, phasic arousal during high probability of rule switches in older people

We used the pupil data to gauge phasic arousal responses to task events as well as latent variables from the best-fitting computational model of the decision process. The pupil dilated robustly during the response trials, with no significant group differences (Fig. 3A), indicating preserved pupil reactivity in older adults. As in previous pupillometry work on change point tasks in younger adults (Nassar et al., 2012; Filipowicz et al., 2020; Murphy et al., 2021; van den Brink et al., 2024), the hidden rule switches also evoked robust pupil responses which peaked around 4 s after the switch (Fig. 3B). This was true for both age groups (Fig. 3B). We compared the pupil response locked with evidence samples where the rule switched with the pupil response locked with a stable rule period (locked to the evidence samples that occurred halfway between two rule switches, with similar temporal statistics as the rule switches). This revealed an effect of condition (rule switch *versus* no rule switch) in two temporal clusters, from –1.5 s up to 0.6 s (smaller response for rule switch; *p* = .002) and from 1.2 s up to 8.5 s (larger response to rule switch; *p* = .001). There was no cluster exhibiting a significant group effect or group x condition interaction.

**Figure 3.**
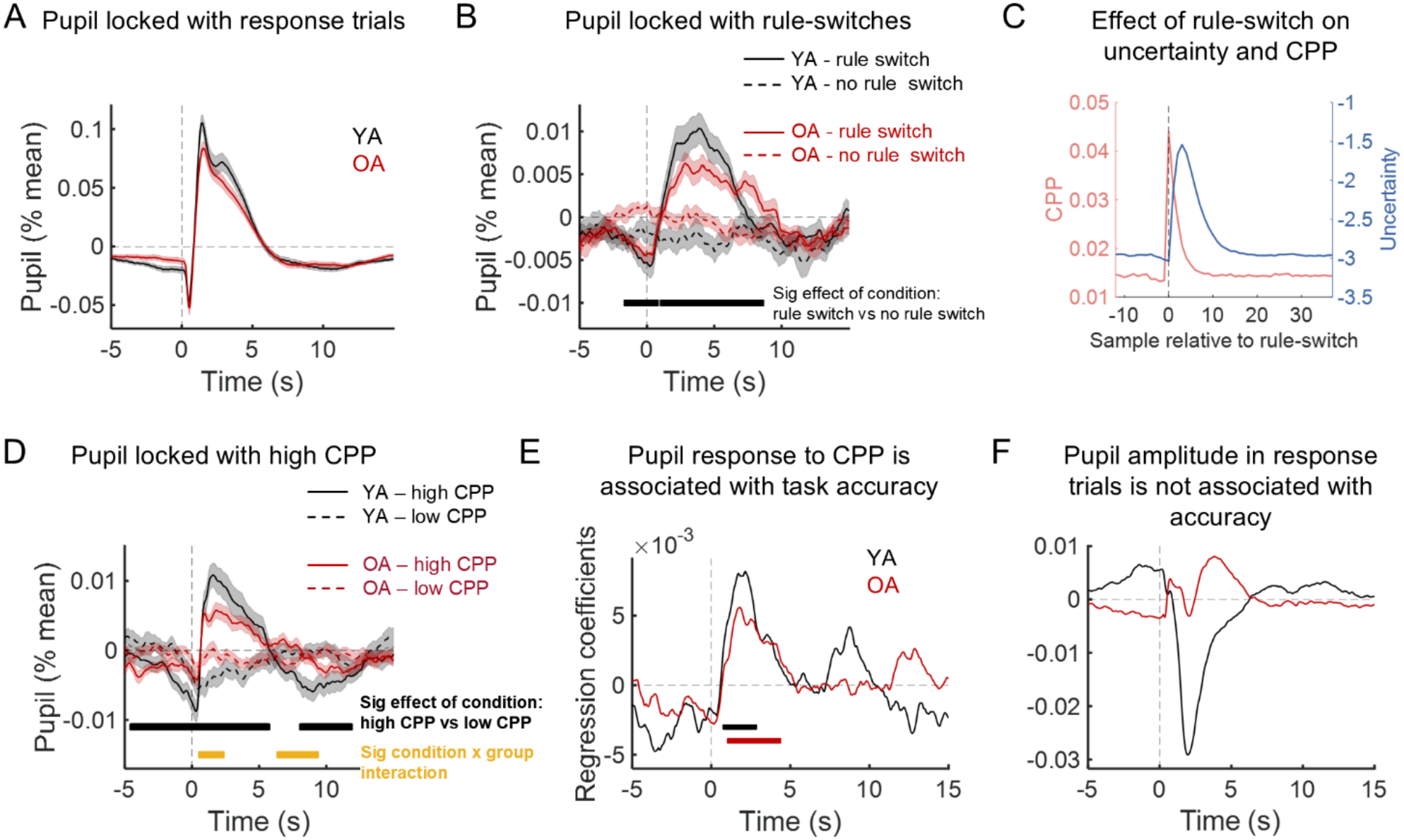
Modulation of pupil-linked arousal by rule switches, model-derived change point probability (CPP), and task accuracy. **A.** Deconvolved pupil response locked with stimulus onset in response trials in the younger (YA) and older adults (OA). **B.** Deconvolved pupil response locked with rule switches (continuous lines) compared with the pupil response locked with the sample halfway between two rule switches (no rule switch; dashed lines). The horizontal black bar marks the window where the response to rule switch was significantly different from the response to no rule switch (repeated measures ANOVA, effect of condition). **C**. Dynamics of model-derived uncertainty and CPP locked with rule switches. **D**. Deconvolved pupil response locked with samples associated with high CPP (continuous lines) or low CPP (dashed lines). The horizontal black bar marks the window where the response to high CPP was significantly different from the response to low CPP (repeated measures ANOVA, effect of condition). The yellow horizontal bar marks the windows where there was a significant group x condition interaction. **E**. Time course of regression coefficients linking task accuracy with the amplitude of pupil response to CPP (calculated as the amplitude difference between the response to high and low CPP). Group specific significant clusters are represented by the black (OA) and red (YA) horizontal lines. **F**. Time course of regression coefficients linking task accuracy with the amplitude of pupil response locked with response trials. In graphs A, B, and D, data is plotted as mean ± standard error of the mean across participants. YA data is plotted in black and OA data in red. All comparisons were corrected for multiple comparisons with a temporal clustering method (Pernet et al., 2015).

How can arousal systems respond to rule switches when those are hidden to the participant? Two latent variables that can be derived from the online inference process described by our behavioral model, increase after rule switches: uncertainty and, particularly sharply, change point probability (CPP) (Fig. 3C; cf, Murphy et al, 2021; van den Brink et al, 2023). Uncertainty reflects (inversely) the strength of the prior belief. CPP is the posterior probability that a rule switch has just occurred, given H, the participant’s current belief, and the new sensory evidence sample, so that the larger the inconsistency between the new evidence sample and the participants’ belief, the larger the CPP. We reasoned that cortical circuits performing the online inference track CPP and uncertainty (McGuire et al., 2014) and drive the brainstem system for pupil control via descending projections, causing the delayed responses to rule switches that we observed. In this case, pupil dynamics should be closely tied to CPP, as observed in other versions of online inference tasks (Nassar et al., 2012; Murphy et al., 2021).

In line with the above idea, samples associated with high CPP evoked stronger pupil responses than samples associated with low CPP in both age groups (Fig. 3D). We found an effect of condition (repeated measures ANOVA, high *vs* low CPP) in three temporal clusters: cluster 1 from –4.4 s up to 0.44 (high CPP < low CPP; *p* = .001), from 0.60 s up to 5.58 s (high CPP > low CPP; *p* = .001) and from 8.22 s up to 11.7 s (high CPP < low CPP; *p* = .003). Critically, the effect of condition depended on age group (significant group x condition interaction), in one cluster starting at 0.70 s and ending at 2.20 s (*p* = .041), and another starting at 6.54 s and ending at 9.20 s (*p* = .006). No significant main effect of group was observed.

The difference between the pupil responses to high and low CPP samples was positively related to individual differences in task performance. We quantified this effect in time point-by-time point regression analyses that revealed a significant association between task accuracy and the pupil high/low CPP amplitude difference in both age groups (younger: one significant temporal cluster starting 0.9 s and ending at 2.7 s, *p* = .036; older: one significant temporal cluster starting 1.2 s and ending at 4.3 s, *p* = .004; Fig. 3E).

We wondered if the association between pupil responses and accuracy was specific to the response during moments of high CPP, or if the more accurate participants showed overall higher pupil reactivity. To this end, we tested the association between the response trial-evoked pupil dilation (Fig. 3A) and accuracy. We found no evidence for an association of these trial-evoked pupil responses and accuracy (Fig. 3F). Moreover, in the younger group, the regression coefficient trended toward negative values, suggesting stronger pupil responses in participants with lower performance. Together, these results argue against a positive association between accuracy and general pupil reactivity.

In sum, there was a transient boost of pupil-linked, phasic arousal during moments of high probability of change in the active rule, which was related to individual task performance. This cognitive drive of the arousal system was weaker in the older group.

### Age effects on pupil responses are not due to task-unrelated factors

Luminance, fatigue, and motivation are all known to affect pupil responses (Lowenstein et al., 1965; Koelewijn et al., 2018; Pan et al., 2022). A number of control analyses showed that these factors were unlikely to explain the smaller pupil responses observed in older participants during high CPP. Luminance pupil responses to the visual stimuli were minimized by design, because all stimuli (photographs and evidence samples) had low contrast, which was identical across all evidence samples and identical for the younger and older adults and therefore were unlikely to explain any differences between age groups. While the onset of the face or house stimuli during response trials caused a small initial pupil constriction, this did not differ between groups (Fig. 3A).

The acquisition site (laboratory *vs* scanner) also did not account for group differences in CPP-evoked pupil responses (Fig. S3). Average pupil size was higher in the scanner than in the laboratory possibly because of the darker environment [*F*_(1, 76)_ = 381, *p* < .001]. However, the difference across acquisition sites was similar across age groups [no group x acquisition site interaction: *F*_(1, 76)_ = 0.12, *p* = .730] and there was no significant group difference [*F*_(1, 76)_ = 0.43, *p* = .512]. Moreover, while the amplitude of the pupil response to high CPP was reduced in the scanner in comparison with the laboratory, this reduction was similar across age groups (no cluster with significant group x acquisition site interaction), and the significant group x condition (high/low CPP) interaction (1 significant cluster between 0.620 s ends at 3.12 s, *p* = .009) was independent of acquisition site (no cluster with significant group x condition x site interaction), suggesting that the group differences were not due to the challenges posed by the MRI environment.

Finally, time-on-task (first half of the session *vs* second half of the session) did not modulate the group effect on CPP-evoked pupil responses (Fig. S4). We found no significant effect of time-on-task or time-on-task x group interaction or 3-way time-on-task x group x condition interaction. These results suggest that the decreased responses observed in the older group were not associated with fatigue and were already present at the beginning of the recordings.

### Relation between task performance and neuropsychological tests

In a final exploratory analysis, we asked which traditional neurocognitive domains assessed in neuropsychological testing are more strongly associated with age-related deficits in our hierarchical decision-making task. Specifically, we evaluated the following cognitive functions based on a comprehensive neuropsychological assessment: processing speed, working memory, and executive control. Performance of our decision task might depend on all these cognitive domains, that are in turn sensitive to age-related decline. For instance, poor processing speed may limit the ability to process the evidence samples presented in rapid succession during online inference (Bruckner et al., 2025), poor working memory might limit the ability to maintain and update evolving belief states (Schapiro et al., 2022; Tavoni et al., 2022), and poor executive control may limit the ability to suppress previously used stimulus-response associations (Miller and Cohen, 2001). We calculated three composite scores each associated with one of the above three cognitive domains (see Methods). Older adults showed significantly lower scores than younger adults in all three domains [independent samples *t*-tests: processing speed *t*_(83)_ = 5.12, *p* < .001; working memory, *t*_(83)_ = 3.72, *p* < .001; executive control *t*_(83)_ = 5.65, *p* < .001].

We then used multiple linear regressions, separately for each age group, to relate the cognitive scores in each domain to accuracy of the hierarchical decision-making task. Variance Inflation Factors for all predictors were below 1.3, indicating no signs of multicollinearity. The overall regression was statistically significant only for older adults (younger adults: *R*^2^ = .168, *F*_(3, 38)_ = 2.35, *p* = .089; older adults: *R*^2^ = .419, *F*_(3, 45)_ = 10.1, *p* < .001), and only the scores for executive control were significantly related to task performance in the older group (Fig. 4). Conversely, when controlling for individual variability in the executive scores, the group difference in accuracy of the hierarchical decision-making task vanished [independent samples *t*-test comparing residual task accuracy after removing effect of executive control scores *t*_(83)_ = 0.808, *p* = .422; Methods]. In other words, reduced performance of older participants in our complex decision-making task was specifically linked to impairments in executive control, and not to other cognitive domains also affected by aging.

**Figure 4.**
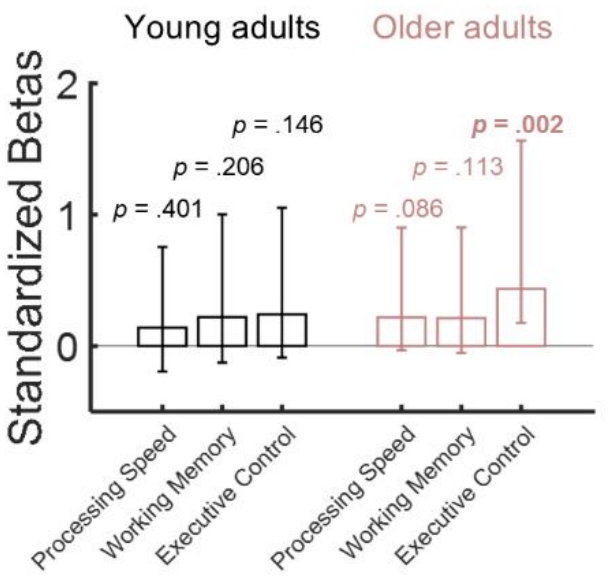
Relationship between performance in the hierarchical decision-making task and in neuropsychological tests assessing processing speed, working memory, and executive control. Multiple linear regressions, separately for each age group, relating the scores in each cognitive domain to the accuracy on the hierarchical decision-making task. Bars represent the standardized regression coefficients for each predictor and error bars represent 95% confidence intervals. Predictors with *p*-values less than .05 are statistically significantly different from zero.

## Discussion

The brainstem arousal systems and in particular the noradrenergic system exhibit changes in structural integrity in healthy and pathological aging which are in turn associated with impairments in memory and executive control (Betts et al., 2017; Liu et al., 2019; Dahl et al., 2020, 2023). The current study investigated whether and how aging affects the transient, top-down activation of these systems during online inference about which context (i.e., stimulus-response mapping rule) for an elementary sensory-motor decision is active, and the rapid adjustment to switches in that context . Here, we approached this question starting from two key insights from previous work in younger adults. Brainstem arousal systems (i) modulate the relative weighting of incoming information during online inference in changing environments (Nassar et al., 2012; O’Reilly et al., 2013; Filipowicz et al., 2020; Murphy et al., 2021, 2024) and (ii) facilitate the rapid remodeling of task-specific sensorimotor networks for flexible behavior (Miller and Cohen, 2001; Fusi et al., 2007; van den Brink et al., 2024). We, therefore, compared the behavior and pupil responses of older and younger individuals performing a hierarchical decision-making task that taps into both online inference as well as the ongoing reconfiguration of sensory-motor associations.

We demonstrate that two findings previously established for younger adults generalize to older adults: older people approximate the normative strategy for solving the hierarchical decision-making task (van den Brink et al., 2023; Calder-Travis et al., 2026) and older people’s pupil-linked arousal is modulated by change point probability in the online inference process included in our task (Nassar et al., 2012; Murphy et al., 2021; van den Brink et al., 2024; Calder-Travis et al., 2026). Older people also seemed to estimate the underlying probability of hidden rule switches equally well as younger adults, detected the changes in hidden rule switches as quickly as younger adults, and also adopted the new rule at the same rate as younger adults.

Critically, we identified one predominant impairment of older compared to younger individuals: a stronger tendency towards using the incorrect stimulus-response mapping rule. This tendency was evident in objective measures (the asymptotic error rates after adjusting to rule switches) as well as participants’ subjective report of rule confusions. The computational model captured this by a higher rate of lapses, random responses, which were independent of the rule evidence. This behavioral impairment was accompanied by smaller pupil responses during moments of high probability of hidden rule switches, which in turn predicted individual task accuracy. Taken together, our findings indicate a link between an age-related abnormal recruitment of brainstem arousal systems and an impaired ability to effectively implement flexible stimulus-response mapping rules. We do not claim that aging selectively impairs the rule implementation . Rather, the latter is the most sensitive aspect of our task. Overall, our results show that in a behavioral context that shares many features with real-life decision situations, the deficient implementation of flexible stimulus-response mapping rules dominates the performance reduction of older compared to younger adults.

The evidence we provide for higher lapse rate in older adults has limitations and alternative scenarios are possible. We asked participants after each block if they realized that they had pressed the wrong button despite knowing the correct decision, and older participants reported a higher number of such perceived errors than younger adults. These subjective reports were obtained after the participants got accuracy feedback, which may have biased their responses. It is possible that this bias differed between age groups. Further, while the lapse parameter in our best-fitting model (and only that parameter) was larger in the older participants, this age effect was weak (statistically significant only before correction for multiple comparisons). Also, while the lapse parameter estimates clearly reflect errors that are independent of the rule evidence, they might reflect other causes of performance fluctuations not captured in the other model parameters (e.g., fluctuations in attention, fatigue, or motivation). Finally, we cannot attribute individual errors in our task to a particular source.

However, several lines of evidence other than subjective report or estimated lapse rates support the idea that rule confusions were the source of errors that most differed between older and younger participants. First, older and younger adults were equally accurate when the lower-order decision (visual categorization) was performed under a fixed stimulus-response mapping rule suggesting no deficit in stimulus categorization. Second, older participants were as accurate as younger participants in inferring the hidden state when they needed to report that directly (“inference-only task”) rather than using it for selecting a rule. Third, after rule switches, accuracy recovered at the same rate in older and younger adults, but only towards a lower asymptotic level indicative of errors during intervals with stable rule.

Our results are overall consistent with some previous work on changes of task switching abilities in aging, which points to no age-related differences in so-called “local” switch costs (the performance costs of changes in rule from one trial to the other are similar across age groups), but significant age-related deficits in “global” switch costs (difference in performance across blocks with rule switches and blocks with stable rules) (Wasylyshyn et al., 2011; Chen and Hsieh, 2023). Performance in our hierarchical decision-making task was also robustly associated with the domain of executive control in neuropsychological assessment, a skill necessary for flexible behavior (Miller and Cohen, 2001). By contrast, task performance was not associated with working memory or processing speed. These findings are compatible with the idea that it is not the inference process, which might depend more on working memory and processing speed, that is limiting older people’s performance but rather poor executive control leading to impulsive random responses associated with rule confusions.

Abnormal recruitment of brainstem arousal systems might limit the neural plasticity necessary for the efficient switch between stimulus-response mapping rules. Both theoretical and experimental (fMRI) work indicate that the flipping of stimulus-response associations in tasks like ours entails the reconfiguration of task-relevant sensory-motor neural pathways through rapid plasticity (van den Brink et al., 2023) governed by brainstem neuromodulatory input (Fusi et al., 2007; van den Brink et al., 2024). It is well established that neuromodulatory (e.g., noradrenergic or serotonergic) systems promote neural plasticity in cortical networks (Bear and Singer, 1986; He et al., 2015; Jordan and Keller, 2023). The activation of brainstem arousal systems at the precise moments when task-relevant networks need to be reconfigured (i.e., high change-point probability) (Bouret and Sara, 2005; Jordan, 2024), might be critical for “imprinting” the new sensory-motor association into the task-relevant neural pathways. The fact that in older adults the arousal responses during moments indicative of rule change is lower might result in impairments in the stabilization of the new sensory-motor association, rendering the rule representation unstable, hindering the efficient implementation of the appropriate stimulus-response association, and leading to higher frequency of rule confusion events. Yet another possibility is that abnormal pupil-linked arousal during moments of high CPP is not causal to the behavioral deficits. With our current correlational approach, we cannot exclude this possibility. Future work could establish causality by transiently boosting neuromodulatory activity (Jordan and Keller, 2023) during rule switching tasks such as ours.

There are several possible explanations for why pupil-linked arousal responses during high CPP are reduced in older people. It is possible that the cognitive recruitment of brainstem arousal systems is impaired in older people due to changes in connectivity limiting their input signals, due to functional limitations of the corresponding brainstem nuclei affecting their output, or both. An alternative possibility is that the observed age-related deficits in pupil responses during moments of high CPP reflect an impaired ability to track that variable (or other latent variables involved in the online inference). Nevertheless, our observation that older participants seemed just as able to perform the inference-only task argues against this account.

In conclusion, in a hierarchical decision-making task, older people showed relatively preserved abilities in inferring hidden changes in behavioral context from noisy evidence but pronounced deficits in the implementation of the context-dependent stimulus-response mapping rules for reporting a lower-order decision. This behavioral deficit co-occurred with alterations in the transient recruitment of pupil-linked arousal systems during moments of high probability of a context/rule switch. Understanding how changes in the brainstem arousal systems affect the ability to switch and implement changeable stimulus-response mappings will be important to design appropriate interventions to ameliorate difficulties in cognitive flexibility in older people.

## Acknowledgments

We would like to acknowledge Hugo Quental for performing the ophthalmologic evaluations of the older participants, Carolina Sousa and Sofia Marcos for their collaboration in the neuropsychological assessment as part of their Curricular Internship and Inês Bernardino and Susana Mouga for their supervision. Finally, we would like to thank all the participants that volunteered for this study for their time and commitment.

