## Supplemental Figures and Tables for "Impaired Sensory-motor Reconfiguration and Pupil-linked Arousal in Aging"

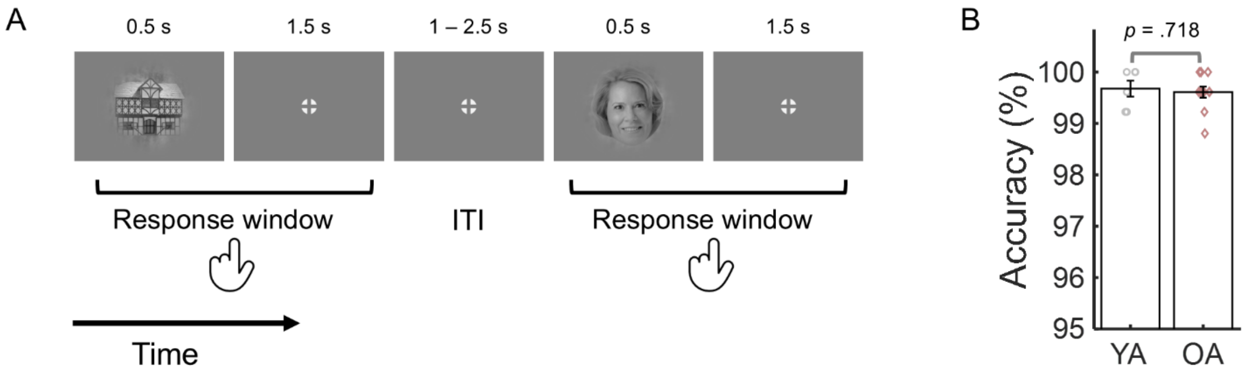


**Figure S1. Categorization performance for the images used in the hierarchical decision-making task. A.** Schematic (not to scale) of the visual categorization task with fixed stimulus-response mapping rule. The same images of faces and houses used in the main task were presented sequentially in random order. Participants were instructed to report if the image was a face or a house through button press with their left or right index fingers. The stimulus-response mapping was fixed for each participant but counterbalanced across participants. **B.** Accuracy as percentage of correct responses for younger (N = 6) and older (N = 11) participants. There was no significant difference across groups in accuracy with all participants responding with high accuracy with a maximum of 3 wrong answers in 258 trials. Two older participants did not respond within the response window in a maximum of 3 trials, and 3 older participants responded with more than one button press in a maximum of 4 trials. Bars represent the group average and error bars represent ± standard error of the mean. Grey circles and red diamonds represent individual data points from younger adults (YA) and older adults (OA), respectively.

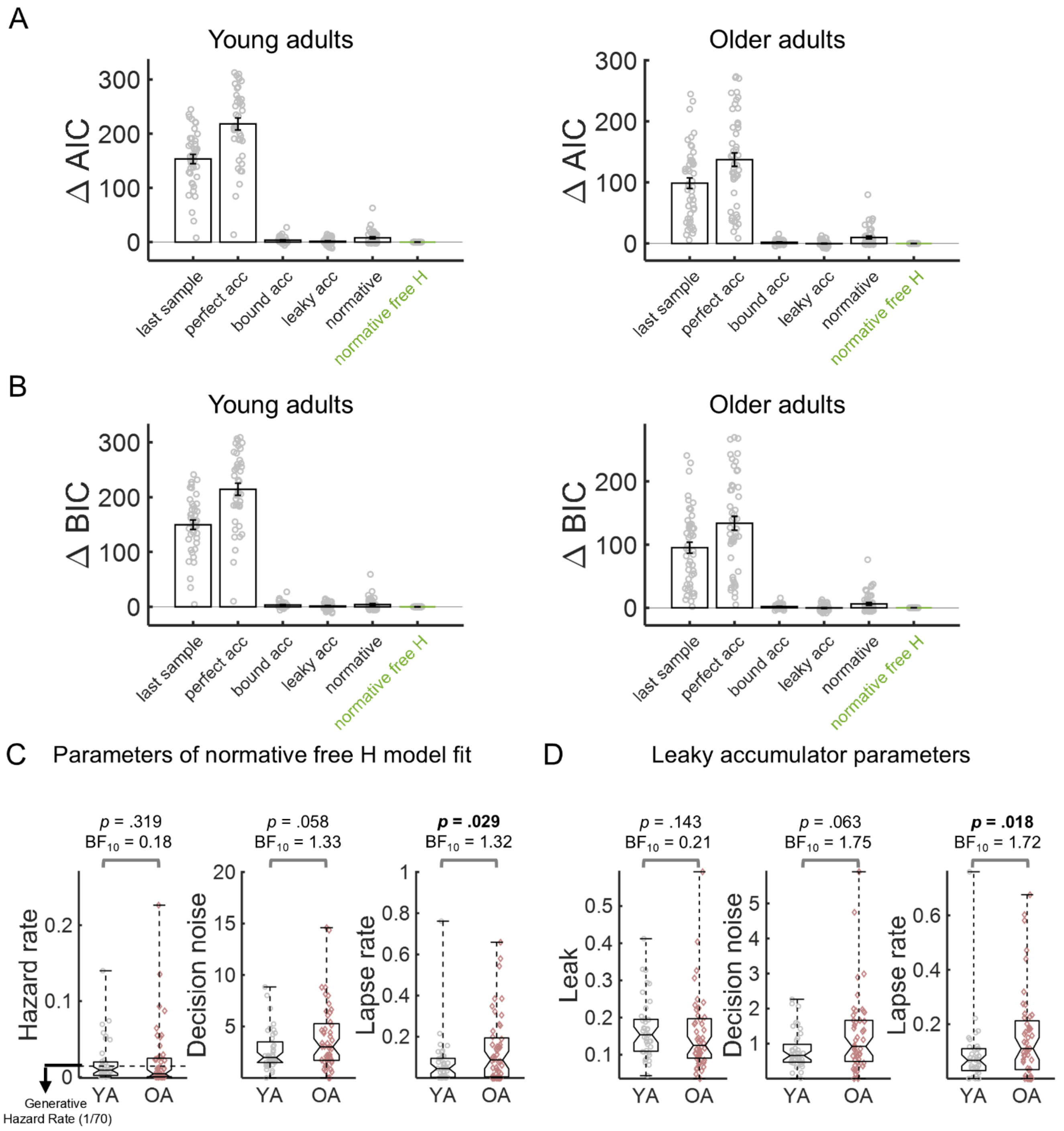


**Figure S2. Alternative models fitted to behavior and fitted model parameters for younger (YA) and older (OA) adults.** **A** and **B**. Model comparison for younger adults (left) and older adults (right) with the normative free H model used as reference (marked in green). All models included decision noise and lapse rate as free parameters. The normative free H model best fitted the behavioral data, according to either AIC or BIC (Methods), except, in the older group, in comparison with the leaky accumulator model with adaptive leak for which the AIC and BIC were slightly lower (ΔAIC = ΔBIC = -0.23). However, these differences were not statistically significant. In fact, for both age groups, the ΔAIC and ΔBIC of all the models (in reference to the normative free H model) were significantly higher than zero (one-sample *t*-tests *p*s < .05) except for the leaky accumulator model (ΔAIC and ΔBIC: younger *p* = .176, older *p* = .743). Note that a leaky accumulator with adaptive leak (Ossmy et al., 2013), approximates the normative model in noise-volatility regimes close to the one used here (Glaze et al., 2015). Grey circles represent individual participants; bars, group average; error bars, ± standard error of the mean. **C.** Parameters from the best fitting normative free H model. Boxplots of parameters of normative free H model fit: hazard rate (left), decision noise (center), and lapse rate (right). Central marks, medians; bottom and top edges, 25th and 75th percentiles; whiskers extend to most extreme data points. *P*-values from Mann–Whitney U test. $BF_{10}$ are Bayes factors for alternative versus the null hypothesis (calculated with Rouder's method assuming unequal variances). Note that the significant difference across groups in lapse rate did not survive adjusting for multiple comparisons (Bonferroni correction, taking into consideration the 3 parameters, lapse rate $p_{adjusted}$ = .087). **D.** Parameters from the best-fitting leaky accumulator model with adaptive leak (Ossmy et al., 2013). OA showed a significant increase in lapse rate, a tendency for higher decision noise and no significant difference for leak parameter in comparison with YA. *P*-values from Mann–Whitney U test. Bayes factors ($BF_{10}$) test the alternative versus the null hypothesis (Rouder's method assuming unequal variance). Note that the significant difference across groups in lapse rate did not survive adjusting for multiple comparisons (Bonferroni correction, taking into consideration the 3 parameters, lapse rate $p_{adjusted}$ = .054). On the boxplots, the central mark indicates the median, and the bottom and top edges of the box indicate the 25th and 75th percentiles, respectively. The whiskers extend to the most extreme data points. Grey circles and red diamonds represent data points from individual young and older participants.

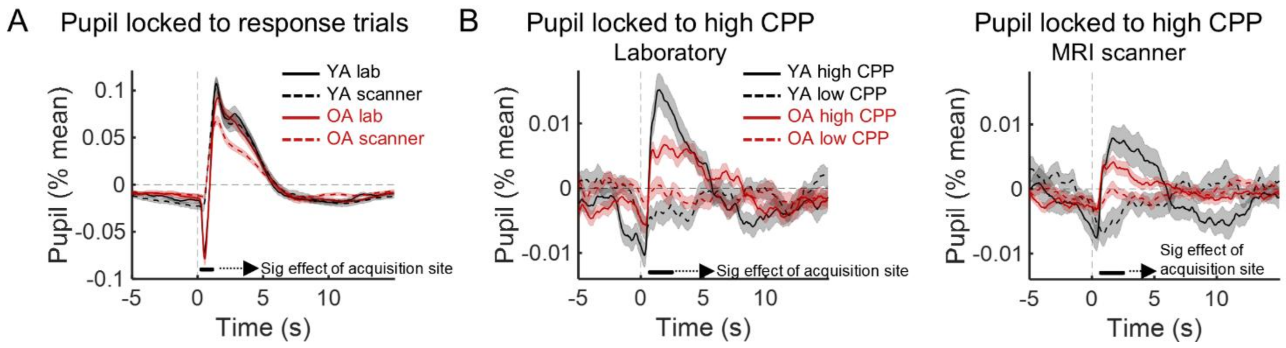


**Figure S3.** Comparison of pupil responses measured in the psychophysics laboratory with the responses measured in the MRI scanner, in younger adults (YA) and older adults (OA), including only the participants with usable eye tracking data from the scanner session (N = 39 YA and N = 39 OA). **A.** Pupil responses locked with the choice trials in the laboratory (continuous lines) and in the scanner (dashed lines). Time point-by-time point repeated measures ANOVA including as within-subject factor acquisition site (lab vs scanner) and as between-subjects factor group revealed a significant effect of acquisition site in the time window marked by the horizontal black line. This window was limited to the period where the pupil showed a luminance response evoked by the visual presentation of the face or house stimuli. This constriction response was more pronounced in the lab than in the scanner due to differences in display systems. There was no significant difference across acquisition sites for the pupil dilation response and no group x site interaction suggesting that the differences in response were similar across groups. **B.** Pupil responses locked with high and low change point probability (CPP) events measured in the laboratory (left) and in the scanner (right). Time point-by-time point repeated measures ANOVA including as within-subject factor acquisition site (lab vs scanner) and condition (high CPP vs low CPP) and as between-subjects factor group, revealed a significant effect of acquisition site with the responses to high CPP showing reduced amplitude in younger and older adults. As in the analyses of the combined lab/scanner data (Fig. 5D), there was a significant effect of condition and a significant condition x group interaction. There was no 3-way interaction condition x group x site suggesting that this effect did not depend on acquisition site. Statistical analyses of the deconvolved pupil responses were performed time point-by-time point using the LIMO EEG toolbox, an open-source MATLAB toolbox (Pernet et al., 2011). We ran second level analysis (group level) using robust trimmed means methods. Results were reported corrected for multiple testing using a temporal clustering forming threshold of p = 0.05 (Pernet et al., 2015).

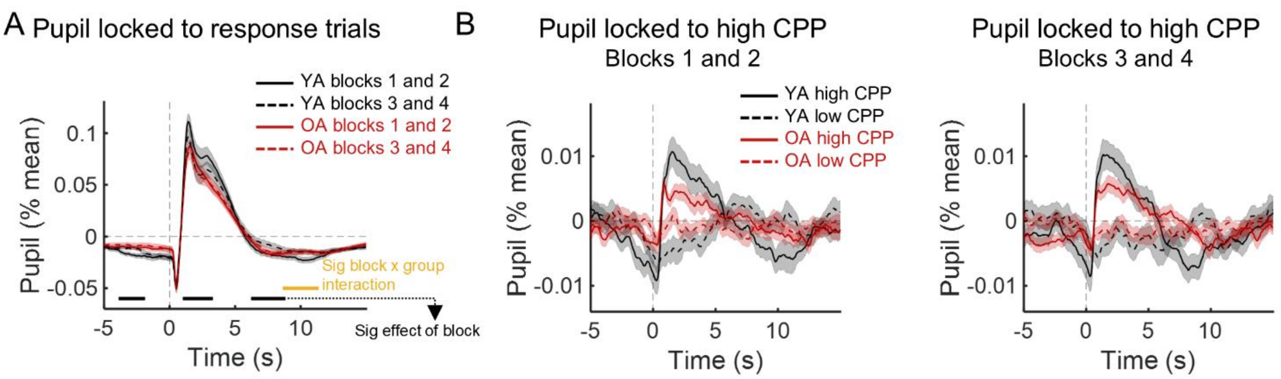


**Figure S4. Study of the effect of time-on-task on pupil responses.** Comparison of pupil responses measured during the first half of the task (blocks 1 and 2 of the laboratory and MRI scanner sessions) and during the second half of the task (blocks 3 and 4 of the laboratory and MRI scanner sessions), in younger adults (YA) and older adults (OA), including only the participants with usable eye tracking data in both periods (N = 40 YA and N = 47 OA). **A.** Pupil responses locked with the response trials in the first half (continuous lines) and in the second half (dashed lines). Time point-by-time point repeated measures ANOVA including as within-subject factor time-on-task (first *vs* second half) and as between-subjects factor group revealed a significant effect of time-on-task in three windows marked by the horizontal black lines reflecting a small reduction of the dilation response in the second half of the task that was similar across groups. A significant time-on-task x group interaction was observed in a later time window around 10 s after the onset of the choice trial that reflected the fact that in that period the pupil response of the younger group was lower in the first in comparison with the second half but this difference was markedly reduced in the older group. **B.** Pupil responses locked with high and low change point probability (CPP) events measured in the first half (left) and in the second half (right) of the task. Time point-by-time point repeated measures ANOVA including as within-subject factor time-on-task (first *vs* second half) and condition (high CPP vs low CPP) and as between-subjects factor group, revealed no significant effect of time-on-task or time-on-task x group interaction or 3-way time-on-task x group x condition interaction. Statistical analyses of the deconvolved pupil responses were performed time point-by-time point using the LIMO EEG toolbox, an open-source MATLAB toolbox (Pernet et al., 2011). We ran second level analysis (group level) using robust trimmed means methods. Results were reported corrected for multiple testing using a temporal clustering forming threshold of $p$ = 0.05 (Pernet et al., 2015).

**Table S1**. Tests, subtests, and measures included in the composite cognitive scores.

| Battery | Cognitive Test | Cognitive Domain | Measure | Composite Score |
|---|---|---|---|---|
| **Wechsler Memory Scale – Third Edition (WMS-III)** | Spatial Span | Working Memory (nonverbal/spatial) | Spatial Span Direct | Working Memory |
| | | | Spatial Span Inverse | Working Memory |
| | Digit Span | Working Memory (verbal) | Digit Span Direct | Working Memory |
| | | | Digit Span Inverse | Working Memory |
| | Letter Number Sequence | Working Memory (verbal) | Letter Number Sequence | Working Memory |
| **CANTAB Connect Research** | Multitasking Test (MTT) | Resistance to Interference | Incongruency cost (median) | – |
| | | | Reaction latency (median) | Processing Speed |
| | | | Multitasking cost (median) | – |
| | | | Total incorrect | Executive Control |
| | One Touch Stockings of Cambridge (OTS) | Planning (visuospatial) | Mean Choices to Correct | Executive Control |
| | | | Problems Solved on First Choice | Executive Control |
| | Reaction Time (RTI) | Processing Speed | Median Five-Choice Reaction Time | Processing Speed |
| | | | Standard Deviation Five-Choice Reaction Time | Processing Speed |

**Table S2**. Factor loadings from a principal-components analysis of the cognitive measures.

| Cognitive Test | Measure | Rotated Factor Loadings[a] | | |
|---|---|---|---|---|
| | | **Factor 1** **Working Memory** | **Factor 2** **Executive Control** | **Factor 3** **Processing Speed** |
| Spatial Span | Spatial Span Direct | **0.687** | -0.343 | 0.051 |
| | Spatial Span Inverse | **0.597** | -0.379 | -0.066 |
| Digit Span | Digit Span Direct | **0.793** | -0.006 | -0.218 |
| | Digit Span Inverse | **0.767** | -0.177 | -0.06 |
| Letter Number Sequence | Letter Number Sequence | **0.74** | -0.076 | -0.315 |
| Multitasking Test (MTT) | Incongruency cost (median) | -0.157 | 0.491 | 0.434 |
| | Reaction latency (median) | -0.392 | 0.456 | **0.61** |
| | Multitasking cost (median) | -0.255 | 0.347 | 0.493 |
| | Total incorrect | -0.261 | **0.619** | -0.047 |
| One Touch Stockings of Cambridge (OTS) | Mean Choices to Correct | -0.1 | **0.882** | 0.176 |
| | Problems Solved on First Choice | 0.141 | **-0.884** | -0.172 |
| Reaction Time (RTI) | Median Five-Choice Reaction Time | -0.155 | -0.013 | **0.792** |
| | Standard Deviation Five-Choice Reaction Time | 0.039 | 0.061 | **0.777** |

[a] Factor loadings are from a principal-components analysis with varimax rotation and Kaiser normalization; loadings of 0.50 or higher are in bold.
